# Social determinants of health, the family, and children's personal hygiene: a comparative study

**DOI:** 10.1101/477943

**Authors:** Antonio Jesús Ramos-Morcillo, Francisco José Moreno-Martínez, Ana María Hernández-Susarte, María Ruzafa-Martínez

## Abstract

**Aims:** To examine differences in personal hygiene and in the perception of social rejection between children in reception centers and children living in a family setting.

**Background:** Little attention has been paid to the influence of the family as a unit on the personal hygiene behaviors of children.

**Design:** Cross-sectional study.

**Methods:** Children aged between 7-12 years were recruited from 2015 through 2017 from two centers in the Network of State Care Centers and from three schools selected from a rural, suburban and urban setting in the same region. A validated questionnaire on child personal hygiene habits was completed by 51 children in reception centers and 454 in normal families.

**Results:** Data shows worse results for the majority personal hygiene habits studied in children in reception centers than in children living in families. Multiple logistic regressions showed lower frequency of body washing, hand washing after defecating, use of soap in hand washing, tooth brushing, and dentist visits during the previous year. Also, a significantly higher proportion of children in reception centers had experienced social rejection for being dirty and smelling bad in comparison to the children living in families.

**Conclusions:** Deficient hygiene habits were observed in the offspring of families affected by the main features of social inequality, who were more likely to perceive social rejection for this reason and less likely to consider their family as the greatest influence on their personal hygiene practices. Promoting family practices designed to improve personal hygiene habits are needed specially in vulnerable families.

## Introduction

Inequalities in health and wellbeing among social groups have been largely attributed to social determinants of health (SDHs), which are considered at least as important as biological mechanisms for disease prevention and treatment [1]. SDHs include socioeconomic and political settings and the particular socioeconomic status of individuals; intermediate determinants of SDHs include material resources, psychosocial, behavioral, and biological factors, and healthcare systems [1].

Multilevel ecological models of SDHs include the family within the “social, family, and community networks” domain, considered not only as a source of support and sustenance but also as an educational resource for the acquisition of healthy habits. Negative aspects are also recognized, with family conflict being a possible risk factor [2]. The family is the first and most important influence on the health and development of children and on the shaping of their routines, habits, attitudes, and social behaviors, including personal hygiene habits [3–5]

## Background

Improvements in sanitary conditions and the acquisition of certain personal hygiene practices during childhood have played a decisive role in reducing infant mortality and increasing life expectancy [6]. However, diseases related to poor hygiene (e.g., diarrhea or respiratory infections) still kill millions of infants in countries with the greatest social inequalities [7–9]. Inadequate hygiene practices have also been implicated in infant morbidity in developed countries, including infectious and parasitic diseases, pneumonia [8], otitis, mycosis, diarrhea, dental caries, gingivitis, and pediculosis [10,11]. Poor hygiene can also be a cause of social rejection, especially for children from poorer families [12].

An inadequate family income is considered as a primary cause of poor health in children [13,14], but the role of the family as social determinant has not been sufficiently considered, although SDH-related factors are known to affect the capacity of families to care for their children [15]. However, researchers have often analyzed the family in a fragmented manner rather than as a unit. For instance, it has been investigated whether the wealth of families and relationships with parents predict healthy behaviors in young people [16] or whether parental educational level is associated with personal hygiene habits [17].

Over recent years, the risk of family poverty has been increased by economic recession, family breakups, and migration, among other factors [18]. Economic inequalities and the lack of effective social policies have also affected the most vulnerable, generating unstructured and dysfunctional families [19]. In extreme cases, such as abuse or abandonment, the state can move children into reception centers for their protection and safety [20]. Children in reception centers (CRCs) have been described as invisible [21], and there has been little research on their health-related lifestyles.

Analysis of the influence of the family as SDH involves the identification of health or healthcare disparity between vulnerable and less-vulnerable populations [22]. The aim of this study was to determine whether CRCs and children living in families (CLFs) differ in their personal hygiene habits and learning and in their perception of social rejection.

## Material and methods

An observational, cross-sectional study compared a group of CRCs with a group of CLFs was carried out.

### Setting and Participants

Children aged between 7 and 12 yrs were studied from March 2015 through January 2017. CRCs were recruited from two centers in the Network of State Care Centers (first stage in fostering process). CLFs were recruited from three schools selected by convenience from a rural (<30,000 inhabitants), suburban (30,000 − 50,000 inhabitants), and urban (>50,000 inhabitants) setting in the same region.

The eligible population was all children in the selected schools and reception centers who met the following inclusion criteria: age between 7 and 12 yrs, voluntary participation, and written consent to participation from parents or legal guardians. Exclusion criteria were inability to speak Spanish or the presence of physical/psychological disabilities that hindered participation. CRCs who had previously been admitted to care were also excluded to avoid the influence of hygiene habits acquired during earlier admission(s).

### Data collection

#### Parents/guardians questionnaire

A questionnaire was administered to parents/guardians of the CLFs to gather data on their educational level, current occupation, type of employment, and household monthly income. CLFs were divided into three groups according to this income: ≤1,000 €, 1,001 - 2,000€, and >2,000 €. Information was also collected on the sex, age, and nationality of the children, the nationality of the parents, number of siblings, and days of school attendance per week.

#### Children Personal Hygiene Questionnaire (HICORIN^®^)

Children data were gathered using the HICORIN^®^ questionnaire, which includes 63 items divided among seven personal hygiene dimensions and hygiene-related social aspects. For this study, we selected items related to the frequency, manner, and timing of personal hygiene activities and the materials used, considering the following dimensions: body skin (8 items), hair (2 items), hands (5 items), oral (14 items); agents affecting personal hygiene learning (8 items); social rejection (2 items); and motivation for personal hygiene activities (5 items).

The HICORIN^®^ questionnaire was interviewer-administered for children aged 7-10 yrs and self-administered for those aged 11-12 yrs. It was completed by participating CLFs at school one week after consent to participation was obtained from parents/guardians and by participating CRCs within two days of admission to the care center. Economic data for the parents/guardians of CRCs were gathered from the computer records of the protection centers.

#### Ethical considerations

The study was approved by the Research Ethics Committee of the University of Murcia (22012014). Written authorization for the children’s participation was obtained from parents/guardians in the case of CLFs and from the General Directorate of Social Policy of Murcia Autonomous Community in the case of CRCs.

#### Data analysis

Exploratory analysis was carried out to evaluate missing data and questionnaires with missing items were eliminated. In a descriptive analysis, we calculated absolute and relative frequencies for qualitative variables and means with standard deviation (SD) for quantitative variables. We used the chi-square test to assess differences in the personal hygiene habits between CRCs and CLFs, stratifying CLFs families into three income levels. We performed multivariate binomial logistic regression analysis to compare the prevalence of hygiene habits between CRCs and CLFs, calculating crude odds ratios (ORs) and ORs adjusted for sex and age with 95 % confidence interval (95% CI). IBM SPSS 21.0 (IBM SPSS, Chicago, IL, USA) was used for the data analysis, considering p<0.05 to be statistically significant.

#### Validity, Reliability and Rigour

HICORIN^®^ questionnaire has been validated for Spanish populations and can be used in healthy children to prevent diseases that might not happen (Moreno-Martínez, Ruzafa-Martínez, Ramos-Morcillo, Gómez García, & Hernández-Susarte, 2015). It has demonstrated adequate reliability; the results of the test-retest coefficient were between very good and moderate in 84.1% of the items. Likewise, content validation, pilot study and items response analysis confirmed satisfactory validity.

## Results

Study eligibility criteria were met by 51 out of the 563 children admitted to the two reception centers during the study period, and all completed the questionnaire (100% response rate). These criteria were met by 758 CLFs in the three schools, and 404 of them completed the questionnaire (53.29% response rate).

### Sociodemographic characteristics of families and children

Comparison of the characteristics of the families of the four study groups (CRCs and three groups of CLFs by monthly household income) revealed significant differences in all study variables (Table 1). In families of CRCs, 90.2% (n=46) of parents had no or only primary schooling, 48% (n=24) were employed (50% of these in unskilled work), and 88.2% (45) of the families had a monthly household income ≤ 1,000€. A significantly higher percentage of CRCs (21 %, n=11) were immigrants in comparison to the three groups of CLFs, 41.2% (n=21) had immigrant mothers and 33% (n=17) had immigrant fathers, 70.6% (n=36) had ≥3 siblings, and 31.4% (n=16) reported not going to school every day.

**Table 1.**
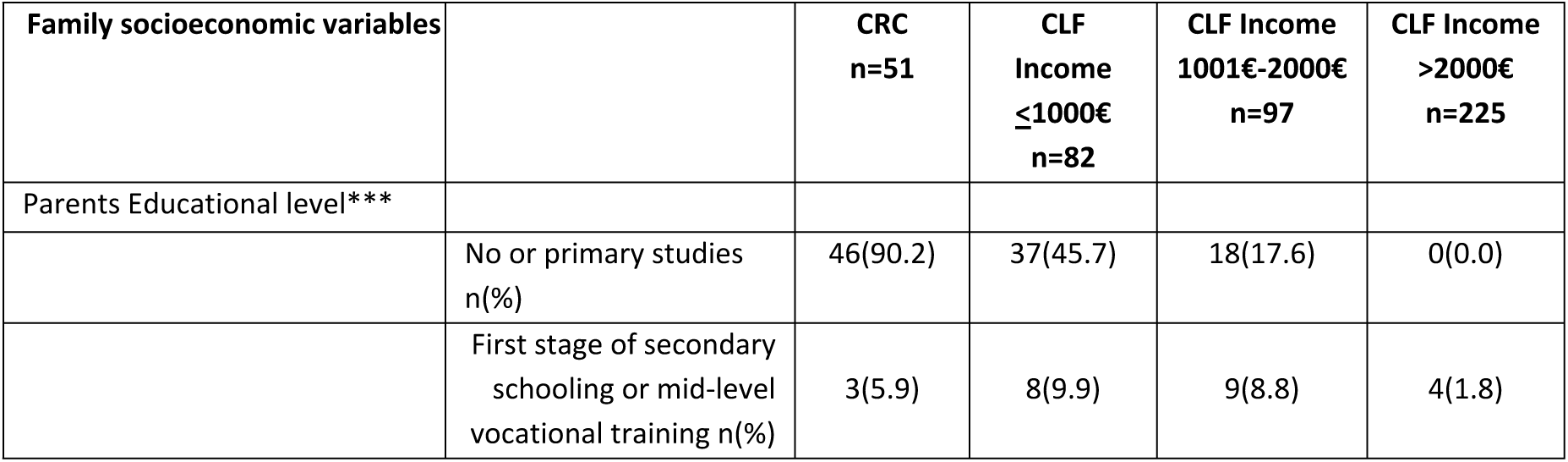
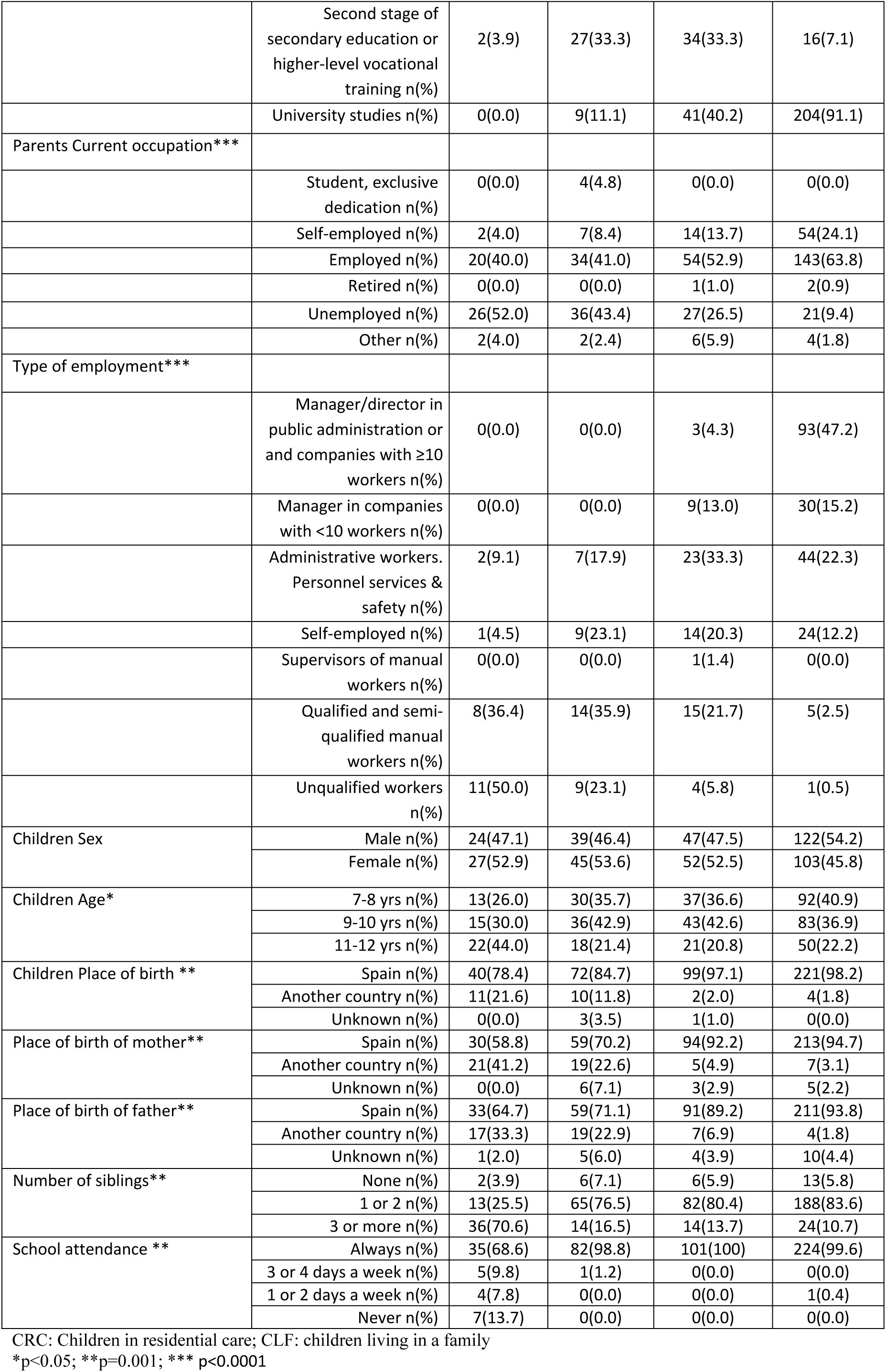
**Comparison of family socioeconomic variables between children in residential care and those living with their family according to its monthly income (N=455)**

| Family socioeconomic variables | | CRC<br>n=51 | CLF<br>Income<br>≤1000€<br>n=82 | CLF Income<br>1001€-2000€<br>n=97 | CLF Income<br>>2000€<br>n=225 |
| --- | --- | --- | --- | --- | --- |
| Parents Educational level*** |  |  |  |  |  |
|  | No or primary studies<br>n(%) | 46(90.2) | 37(45.7) | 18(17.6) | 0(0.0) |
|  | First stage of secondary<br>schooling or mid-level<br>vocational training n(%) | 3(5.9) | 8(9.9) | 9(8.8) | 4(1.8) |
|  | Second stage of secondary education or higher-level vocational training n(%) | 2(3.9) | 27(33.3) | 34(33.3) | 16(7.1) |
|  | University studies n(%) | 0(0.0) | 9(11.1) | 41(40.2) | 204(91.1) |
| Parents Current occupation*** |  |  |  |  |  |
|  | Student, exclusive dedication n(%) | 0(0.0) | 4(4.8) | 0(0.0) | 0(0.0) |
|  | Self-employed n(%) | 2(4.0) | 7(8.4) | 14(13.7) | 54(24.1) |
|  | Employed n(%) | 20(40.0) | 34(41.0) | 54(52.9) | 143(63.8) |
|  | Retired n(%) | 0(0.0) | 0(0.0) | 1(1.0) | 2(0.9) |
|  | Unemployed n(%) | 26(52.0) | 36(43.4) | 27(26.5) | 21(9.4) |
|  | Other n(%) | 2(4.0) | 2(2.4) | 6(5.9) | 4(1.8) |
| Type of employment*** |  |  |  |  |  |
|  | Manager/director in public administration or and companies with ≥10 workers n(%) | 0(0.0) | 0(0.0) | 3(4.3) | 93(47.2) |
|  | Manager in companies with <10 workers n(%) | 0(0.0) | 0(0.0) | 9(13.0) | 30(15.2) |
|  | Administrative workers. Personnel services & safety n(%) | 2(9.1) | 7(17.9) | 23(33.3) | 44(22.3) |
|  | Self-employed n(%) | 1(4.5) | 9(23.1) | 14(20.3) | 24(12.2) |
|  | Supervisors of manual workers n(%) | 0(0.0) | 0(0.0) | 1(1.4) | 0(0.0) |
|  | Qualified and semi-qualified manual workers n(%) | 8(36.4) | 14(35.9) | 15(21.7) | 5(2.5) |
|  | Unqualified workers n(%) | 11(50.0) | 9(23.1) | 4(5.8) | 1(0.5) |
| Children Sex | Male n(%) | 24(47.1) | 39(46.4) | 47(47.5) | 122(54.2) |
|  | Female n(%) | 27(52.9) | 45(53.6) | 52(52.5) | 103(45.8) |
| Children Age* | 7-8 yrs n(%) | 13(26.0) | 30(35.7) | 37(36.6) | 92(40.9) |
|  | 9-10 yrs n(%) | 15(30.0) | 36(42.9) | 43(42.6) | 83(36.9) |
|  | 11-12 yrs n(%) | 22(44.0) | 18(21.4) | 21(20.8) | 50(22.2) |
| Children Place of birth ** | Spain n(%) | 40(78.4) | 72(84.7) | 99(97.1) | 221(98.2) |
|  | Another country n(%) | 11(21.6) | 10(11.8) | 2(2.0) | 4(1.8) |
|  | Unknown n(%) | 0(0.0) | 3(3.5) | 1(1.0) | 0(0.0) |
| Place of birth of mother** | Spain n(%) | 30(58.8) | 59(70.2) | 94(92.2) | 213(94.7) |
|  | Another country n(%) | 21(41.2) | 19(22.6) | 5(4.9) | 7(3.1) |
|  | Unknown n(%) | 0(0.0) | 6(7.1) | 3(2.9) | 5(2.2) |
| Place of birth of father** | Spain n(%) | 33(64.7) | 59(71.1) | 91(89.2) | 211(93.8) |
|  | Another country n(%) | 17(33.3) | 19(22.9) | 7(6.9) | 4(1.8) |
|  | Unknown n(%) | 1(2.0) | 5(6.0) | 4(3.9) | 10(4.4) |
| Number of siblings** | None n(%) | 2(3.9) | 6(7.1) | 6(5.9) | 13(5.8) |
|  | 1 or 2 n(%) | 13(25.5) | 65(76.5) | 82(80.4) | 188(83.6) |
|  | 3 or more n(%) | 36(70.6) | 14(16.5) | 14(13.7) | 24(10.7) |
| School attendance ** | Always n(%) | 35(68.6) | 82(98.8) | 101(100) | 224(99.6) |
|  | 3 or 4 days a week n(%) | 5(9.8) | 1(1.2) | 0(0.0) | 0(0.0) |
|  | 1 or 2 days a week n(%) | 4(7.8) | 0(0.0) | 0(0.0) | 1(0.4) |
|  | Never n(%) | 7(13.7) | 0(0.0) | 0(0.0) | 0(0.0) |

CRC: Children in residential care; CLF: children living in a family
\*p<0.05; \*\*p=0.001; \*\*\* p<0.0001

### Personal hygiene habits

Tables 2 exhibits the results for body and hair hygiene habits. Statistically significant differences were found between CRCs and CLFs in almost all items. The frequency of body washing “≥3 days a week” was 12- to 15-fold lower in CRCs than in CLFs; a wet towel/sponge was used for body washing by 23.5% (n=12) of CRCs *versus* almost 100% of CLFs who reported taking a shower/bath; and body washing was performed at night (before bedtime) by 39.2% (n=20) of CRCs *versus* <50 % of CLFs. With regard to materials, gels were not used or known by 21.6% (n=11) of CRCs *versus* 3.6-7.1% of CLFs, the use of bars of soap was uncommon but was more frequent by CRCs than by CLFs (n=17). A washbowl was more often used in body washing by CRCs than by CLFs, whose use of this complement was less frequent with higher household income. A hair washing frequency of “≥3 times/week” was 8.6-9.5-fold lower in CRCs than in the CLF groups. The frequency of shampoo use did not significantly differ between CRCs and CLFs (p=0.364).

**Table 2.**
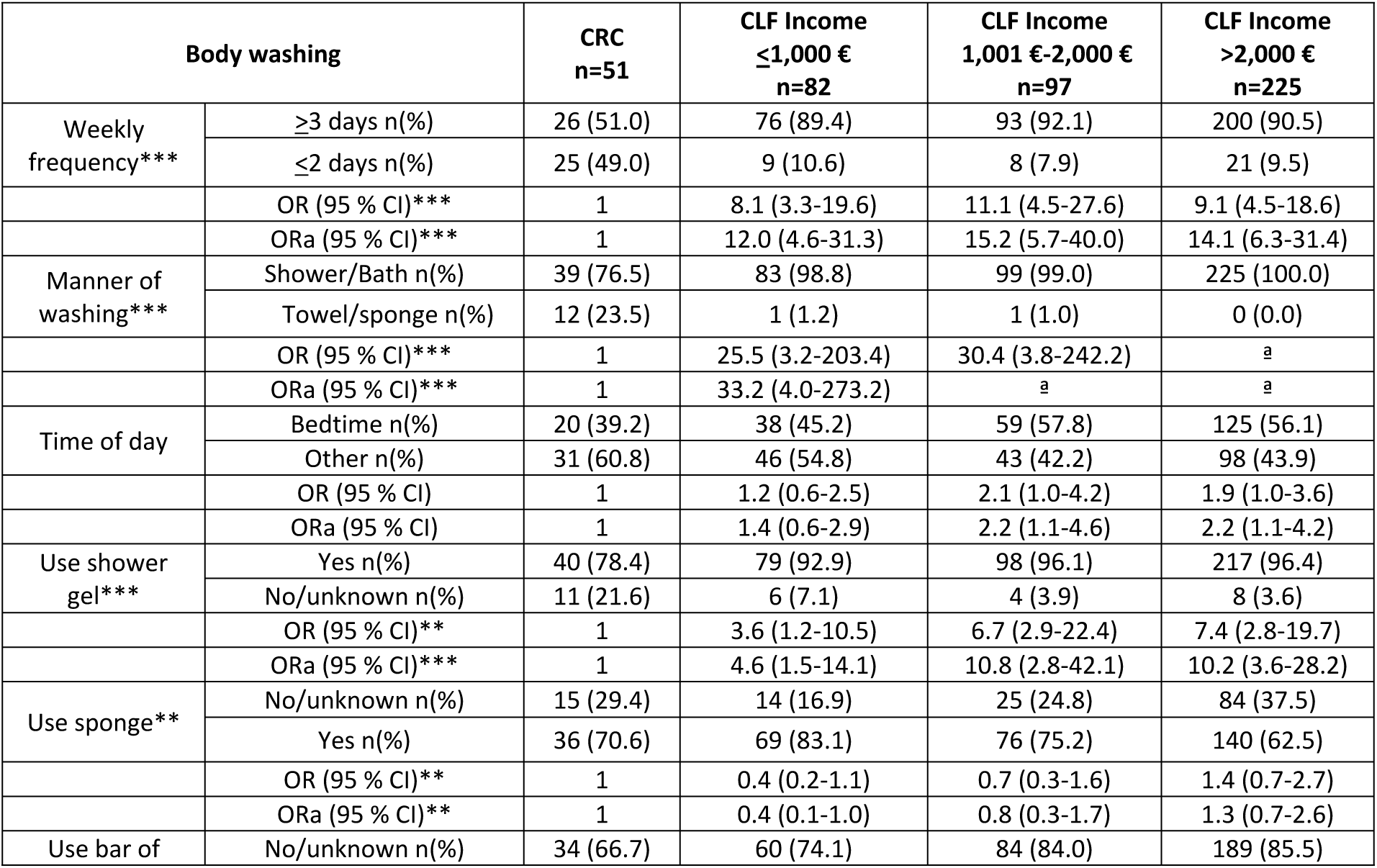
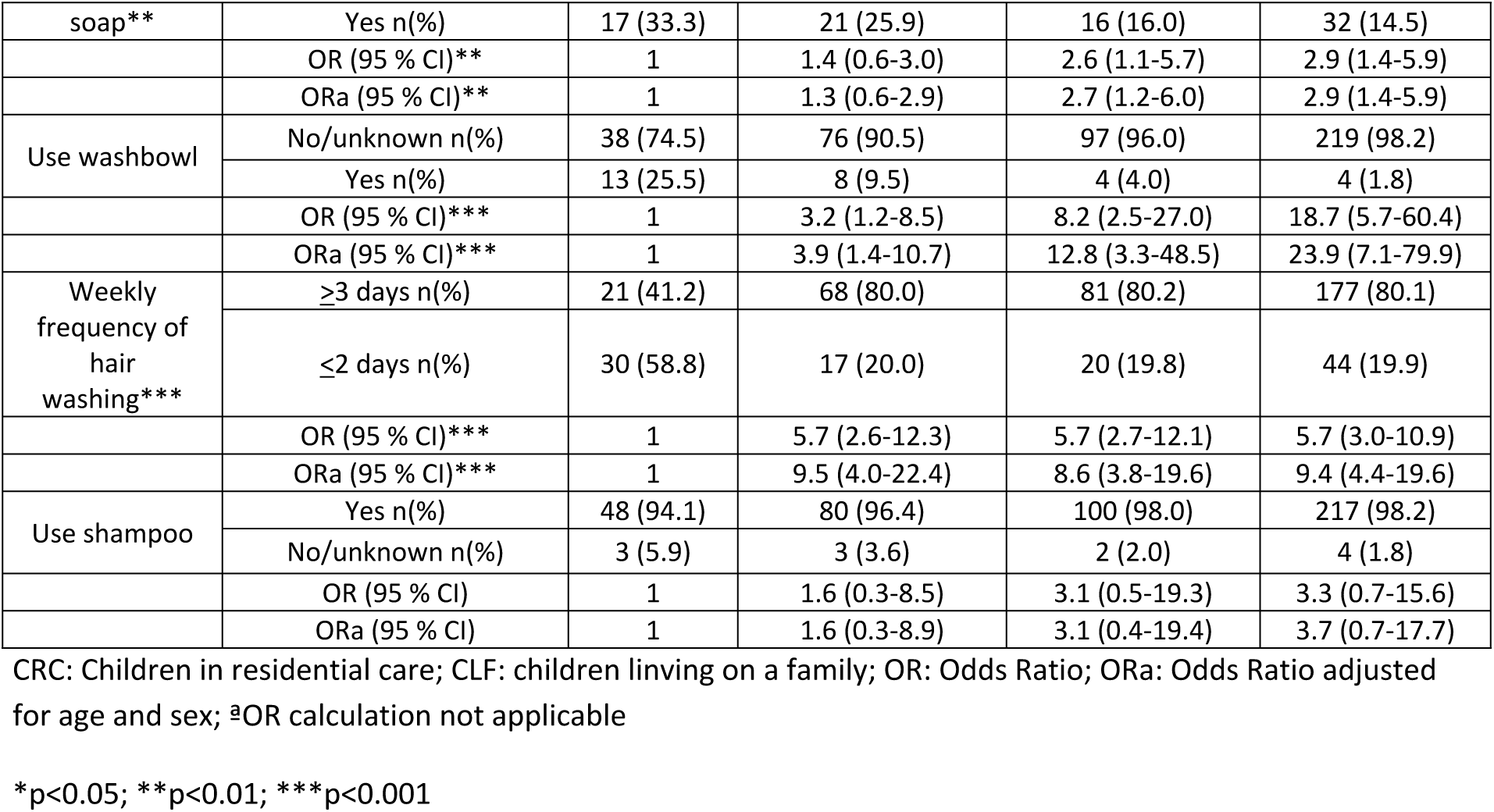
**Comparison of body/hair and hand hygiene variables between children in residential care and those living with their family according to its monthly income (N=455)**

As shown in Table 3 “hand washing ≥3times a day” was reported by a lower percentage of CRCs than of CLFs, regardless of their household income, but the difference was not statistically significant. The use of soap in hand washing was significantly less frequent among CRCs (39.2%; 20) than among CLFs in the income group between 1,001 and 2,000€ (89.2%; 91). Statistically significant differences were also obtained in hand washing frequencies, which were always lower for CRCs. Hand washing after defecating was the most frequent practice in all study groups, although it was reported by a higher percentage of CLFs than CRCs.

**Table 3.** **Comparison of hand hygiene variables between children in residential care and those living with their family according to its monthly income (N=455)**

| Hand washing |  | CRC<br>n=51 | CLF<br>Income ≤1,000 €<br>n=83 | CLF Income<br>1,001 €-2,000 €<br>n=97 | CLF<br>Income >2,000 €<br>n=224 |
| --- | --- | --- | --- | --- | --- |
| Daily frequency | >3 times n(%) | 13 (25.5) | 31 (36.5) | 35 (35.0) | 80 (35.7%) |
|  | ≤3 times n(%) | 38 (74.5) | 54 (63.5) | 65 (65.0) | 144 (64.3) |
|  | OR (95 % CI) | 1 | 1.6 (0.7-3.6) | 1.5 (0.7-3.3) | 1.6 (0.8-3.2) |
|  | ORa (95 % CI) | 1 | 2.0 (0.9-4.4) | 1.7 (0.8-3.8) | 1.9 (0.9-3.9) |
| Always before every meal* | Yes n(%) | 5 (9.8) | 47 (55.3) | 53 (52.0) | 116 (51.6) |
|  | No n(%) | 46 (90.2) | 38 (44.7) | 49 (48.0) | 109 (48.4) |
|  | OR (95 % CI)* | 1 | 11.3 (4.1-31.4) | 9.9 (3.6-27.0) | 9.7 (3.7-25.5) |
|  | ORa (95 % CI)* | 1 | 12.0 (4.3-33.3) | 9.9 (3.6-27.2) | 9.7 (3.7-25.5) |
| Always after | Yes n(%) | 16 (31.4) | 60 (70.6) | 79 (77.5) | 172 (77.1) |
| defecating* | No n(%) | 35 (68.6) | 25 (29.4) | 23 (22.5) | 51 (22.9) |
|  | OR (95 % CI)* | 1 | 5.2 (2.4-11.1) | 7.5 (3.5-15.9) | 7.3 (3.7-14.4) |
|  | ORa (95 % CI)* | 1 | 9.5 (4.1-22.1) | 11.2 (4.9-25.6) | 12.8 (6.0-27.5) |
| Always after urinating* | Yes n(%) | 7 (13.7) | 47 (55.3) | 61 (59.8) | 121 (53.8) |
|  | No n(%) | 44 (86.3) | 38 (44.7) | 41 (40.2) | 104 (46.2) |
|  | OR (95 % CI)* | 1 | 7.7 (3.1-19.2) | 9.3 (3.8-22.7) | 7.3 (3.1-16.9) |
|  | ORa (95 % CI)* | 1 | 10.6 (4.1-27.4) | 11.9 (4.7-30.2) | 10.1 (4.2-24.3) |
| Always use soap* | Yes n(%) | 20 (39.2) | 63 (74.1) | 91 (89.2) | 181 (80.4) |
|  | No n(%) | 31 (60.8) | 22 (25.9) | 11 (10.8) | 44 (19.6) |
|  | OR (95 % CI)* | 1 | 4.4 (2.1-9.3) | 12.8 (5.5-29.7) | 6.3 (3.3-12.2) |
|  | ORa (95 % CI)* | 1 | 4.5 (2.1-9.6) | 13.6 (5.7-32.3) | 6.3 (3.3-12.2) |

CRC: children in residential care; CLF: Children living on a family; OR: Odds Ratio; ORa: Odds Ratio adjusted for age and sex
\*p<0.001

Statistically significant results were observed in all oral hygiene items (Table 4). The tooth brushing frequency “≥2times a day” was reported by a lower percentage of CRCs than of CLFs, and the percentage of CLFs increased with higher household incomes. The highest frequency of tooth brushing by all of the children was at night (before bedtime), although it was significantly less frequent in CRCs. With respect to the materials used for oral hygiene, toothbrush and toothpaste were the most frequently used (>80% in all groups). In addition, 17.6% (9) of CRCs shared their toothbrush with other family members. There were also significant differences in the type of toothbrush used, with 6% of CRCs reporting the use of an electrical toothbrush *versus* 40% of CLFs. Conversely, utilization of a toothpick was reported by a higher percentage of CRCs than of CLFs (37.3%; 19). The frequency of dentist visits during the previous year was significantly lower in CRCs and was greater with higher household income in CLFs.

**Table 4.**
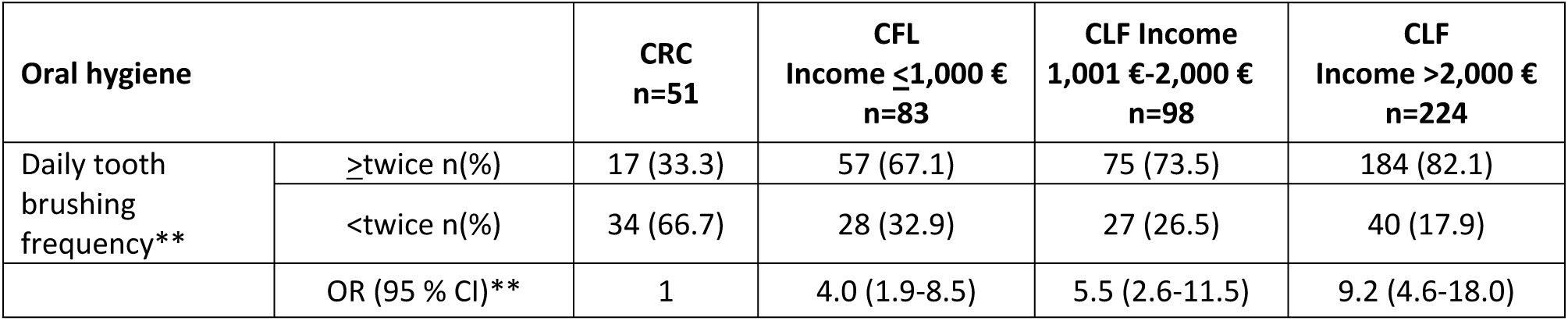
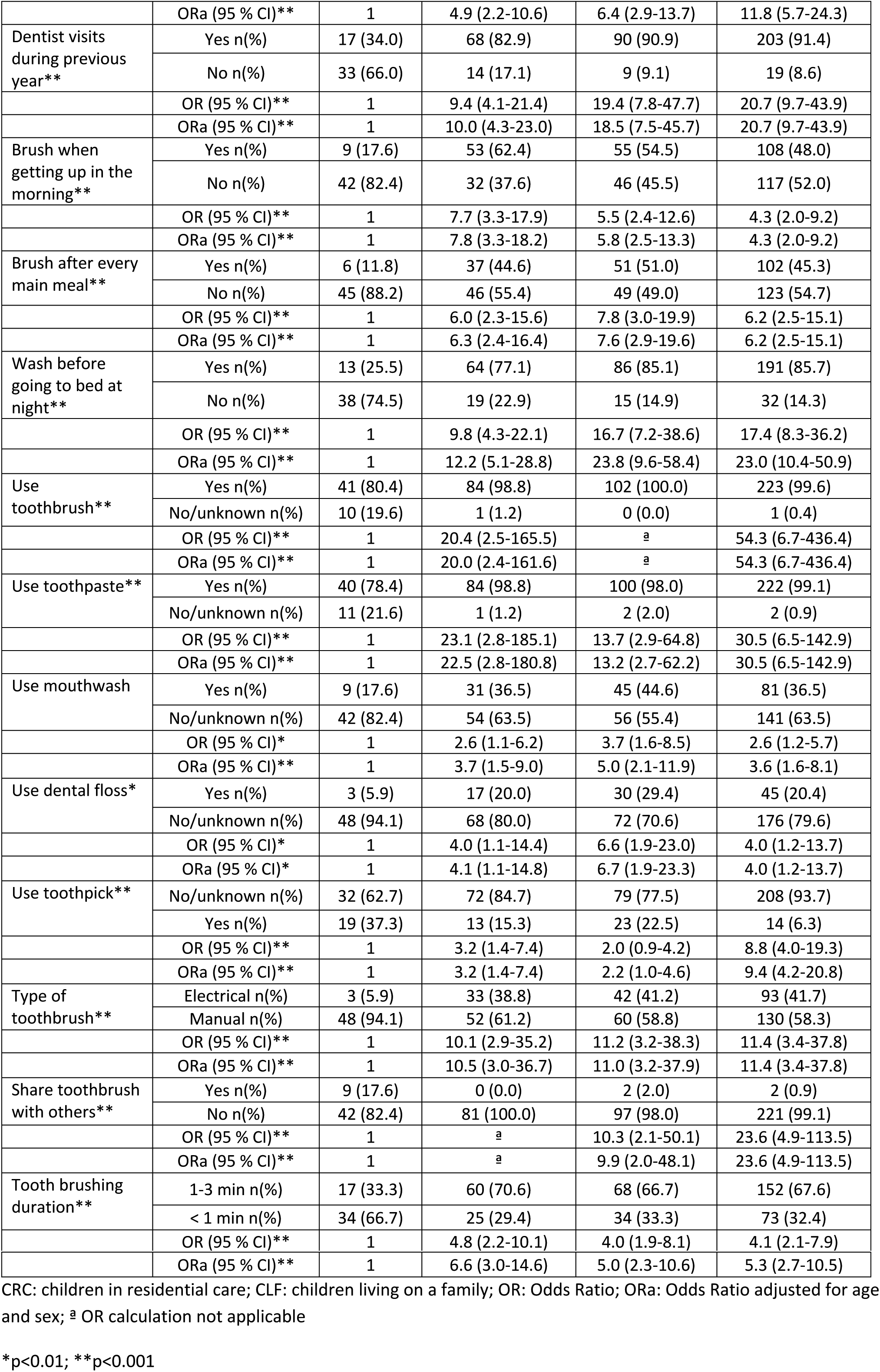
**Comparison of oral hygiene variables between children in residential care and those living with their family according to its monthly income (N=456)**

### Personal hygiene practice learning and social rejection

The family (father, mother, or other family member) was most frequently described by all groups of children as having the greatest influence on their personal hygiene learning, although this affirmation was made by a significantly lower percentage of CRCs (72.5 %, 37) than of CLFs. In second place as learning agents, CLFs selected healthcare professionals, while CRCs selected teachers, radio, television, internet, and self-learning (Table 5).

A significantly higher proportion of CRCs (41.1 %) had experienced social rejection for being dirty in comparison to the CLFs (9.5% of those with family incomes< 1.000€ and around 4% of those with higher family incomes). Very similar differences were observed in the experience of rejection for smelling bad, although the percentages were slightly higher in all groups, reaching almost 50% in CRCs and between 8.9 and 10.7%in CLFs. The only statistically significant difference in motivations for personal hygiene activities was in the option “to not be rejected by friends”, which was selected by 90% of CRCs *versus* 58% of CLFs from families with incomes < 1,000€ and 36% of those from families with higher incomes.

**Table 5.**
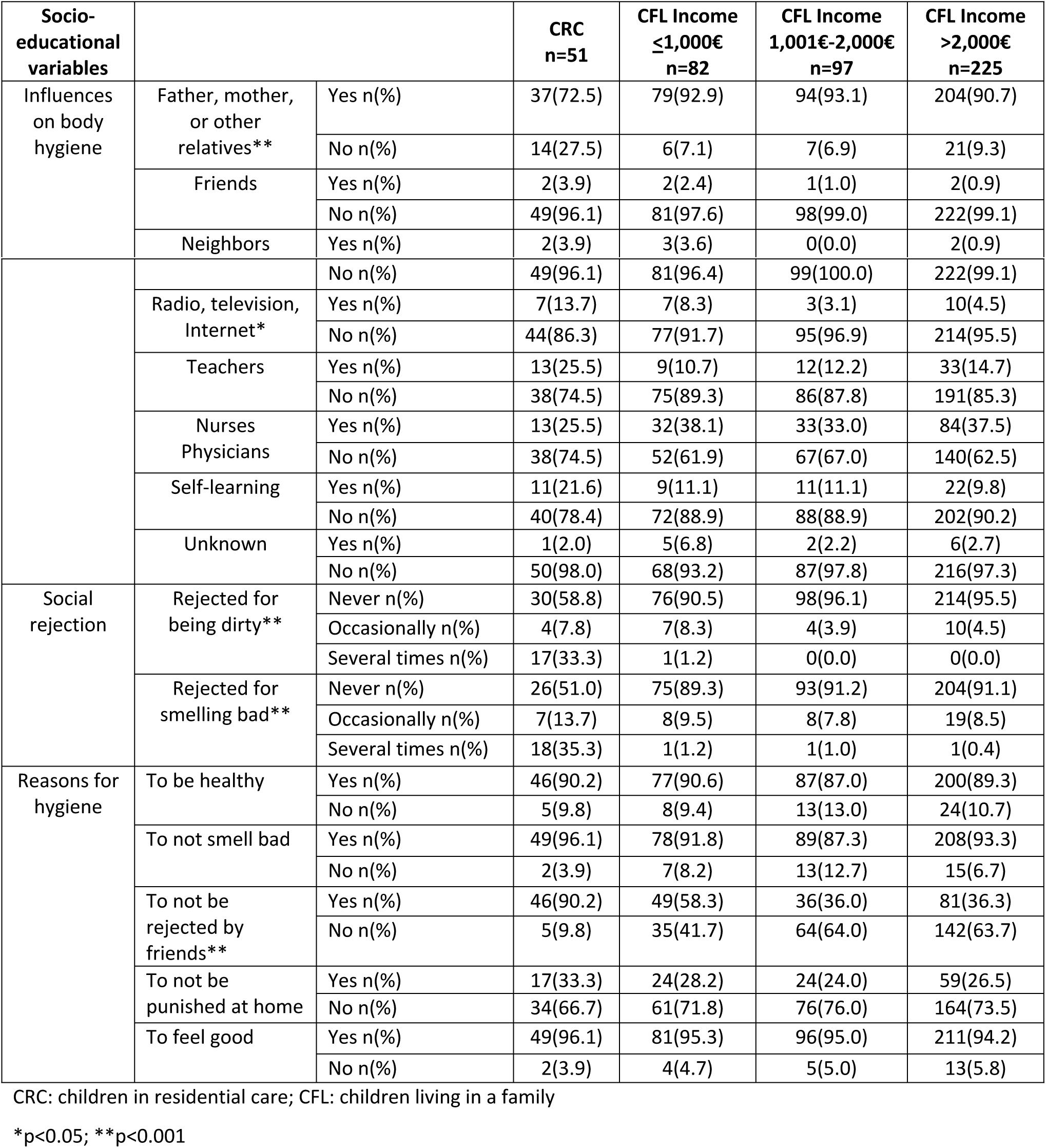
**Comparison of socio-educational variables between children in residential care and those living with their family according to its monthly income (N=455)**

## Discussion

There has been little research in developed countries on SDHs related to family and personal hygiene in childhood, and this study therefore contributes important empirical data. The main finding was a clear relationship between the personal hygiene of the children and their family settings.

In this study, CLFs were compared with CRCs whose situation of vulnerability was sufficiently extreme to warrant removal from their families [23]. The parents/guardians of the CRCs exhibit the main axes of inequality [1], being characterized by a low schooling level, low qualifications, unemployment or only casual unskilled employment, and an income < 1,000€, with 70% having ≥ 3 children. Among the CRCs, 78.4% were born in Spain, although around two out of five of their parents were immigrants, and almost one out of three children did not go to school every day.

In comparison to the CLFs, the CRCs had poorer hygiene habits in all dimensions studied, were less likely to consider their families as the most influential agent in learning hygiene habits, and were more likely to experience social rejection due to their hygiene and to be motivated by this rejection to carry out personal hygiene activities. These findings question whether vulnerable families are adequately fulfilling their functions of protection, healthcare, and socialization [22,24].

The lower frequency of key personal hygiene practices (body, hair hand washing and tooth brushing) in the children from vulnerable families is consistent with reports that implicate the family structure and low parental educational and socioeconomic level in poor health, hygiene [25], and oral hygiene [26] behaviors in children. A strong relationship has been reported between low level of healthy habits and worse children’s health as perceived by their mothers (OR=0.48 95% CI=0.42-0.56) [25]. Hence, the detection of signs of poor personal hygiene practices in children may help professionals to anticipate situations of ill health.

As previously observed by [27], an association was found between lower family income and worse hygiene practices, especially dental hygiene habits, with less frequent daily tooth brushing, tooth brushing before bedtime, and visits to the dentist. Despite the offer of free oral healthcare, visits to the dentist during the previous year were between 10- and 20.7-fold less frequent for CRCs than for CLFs. Researchers have concluded that families with lesser resources have a worse relationship with healthcare systems [28,29], which they engage with to a lesser extent, in part due to the lack of recognition of health problems [30].

The relationship between the physical environment in which children develop and healthy behaviors has also been demonstrated [3–5,31]. In the present study, children of vulnerable families were less likely to use a toothbrush (especially an electrical device), hand-washing soap, shower gel, or toothpaste, while one in four of them used a washbowl and wet towel or sponge for body washing, reflecting the lack of opportunity to take a bath or shower.

Surprisingly, hand washing was not an established routine in the children, regardless of their family or institutional setting, possibly because it was carried out without the supervision of parents, despite being recommended as a family activity [32]. There was a higher frequency of hand washing after defecation in both CRCs and CLFs, although it continued to be low in the former, similar to the findings of a study in 11 developing countries [33]. This practice requires special attention, given that education in hand washing has been reported to reduce cases of diarrhea by 31% and cases of respiratory diseases by 21% [7,34].

Besides the biological implications of our findings, they are also relevant from a social standpoint, confirming the value of the family as a key factor in the acquisition of healthy habits [35]. Almost all CLFs and three-quarters of CRCs considered their families to be the main agents for learning hygiene practices, with a role also being attributed by CLFs to healthcare professionals and by CRCs to teachers, self-learning, and the mass media, which may reflect the worse relationship of poorer families with healthcare systems [28,29]. These findings suggest the need for educators and healthcare professionals to work together in the design and implementation of strategies to improve the hygiene habits of children from vulnerable families.

Weaknesses in the socialization function of the family were also revealed by the CRCs, who were more likely to be rejected for being dirty or smelling bad and to be motivated in their personal hygiene activities by the need to avoid this rejection. Hygiene behaviors play an important role in the impression that we make on others and as a display of respect for social norms [12], facilitating the integration and socialization of children among their peers. There is a relationship among family vulnerability, diseases associated with poor hygiene (e.g., caries and pediculosis), school problems, and marginalization [17,36,37]. The long-term social effects of this situation are unclear; however, it appears likely to perpetuate the inequality and vulnerability of these children, with negative consequences for their health in adulthood [38]. According to the Model of Health Promotion of the Family [35], the values, targets, and needs of families mediate between their health practices and socioeconomic status. Hence, despite the limited influence of nursing professionals on the SDHs affecting families, they may be able to intervene in these mediators to improve health and hygiene practices. Family interventions have achieved improvements in the acquisition of healthy lifestyles related to physical activity and sport [39] and might therefore have a similar impact on personal hygiene habits.

### Limitations

This study contains certain limitations. The sample size of children from vulnerable families was reduced due to difficulties in gaining access to this relatively small population, although it proved possible to obtain significant differences with the children living at home. There may have been a “volunteer” bias, given the response rate of only 60% for the CLFs, and the study design means that causality relationships could not be established. Finally, the lack of published data on this issue, especially in developed countries, limited the discussion of our findings.

## Conclusions

Our study provides new evidence on the relationship among SDHs, family, and the personal hygiene practices of children. Our findings raise questions about the adequate fulfillment by vulnerable families of their protection, healthcare, and socialization functions. The results confirm that the family, understood as a complex system that acts on the health behaviors of the individuals that form it, affects the personal hygiene practices of children. Thus, deficient hygiene habits were observed in the offspring of families affected by the main features of social inequality, who were more likely to perceive social rejection for this reason and less likely to consider their family as the greatest influence on their personal hygiene practices.

These findings indicate that action against social inequality can have a potential impact on biological mechanisms that affect health. Although this inequality cannot be resolved within the family setting, it can be ameliorated by promoting family practices designed to improve personal hygiene habits.

## Acknowledgements

The authors are grateful to all of the children, families, schools and reception centers and their staff, and to the General Directorate of Social Policy of Murcia Autonomous Community.

